# Look where you go: characterizing eye movements toward optic flow

**DOI:** 10.1101/2020.10.02.324384

**Authors:** Hiu Mei Chow, Jonas Knöll, Matthew Madsen, Miriam Spering

## Abstract

When we move through our environment, objects in the visual scene create optic flow patterns on the retina. Even though optic flow is ubiquitous in everyday life, it is not well understood how our eyes naturally respond to it. In small groups of human and non-human primates, optic flow triggers intuitive, uninstructed eye movements to the pattern’s focus of expansion (Knöll, Pillow & Huk, 2018). Here we investigate whether such intuitive oculomotor responses to optic flow are generalizable to a larger group of human observers, and how eye movements are affected by motion signal strength and task instructions. Observers (*n* = 43) viewed expanding or contracting optic flow constructed by a cloud of moving dots radiating from or converging toward a focus of expansion that could randomly shift. Results show that 84% of observers tracked the focus of expansion with their eyes without being explicitly instructed to track. Intuitive tracking was tuned to motion signal strength: saccades landed closer to the focus of expansion and smooth tracking was more accurate when dot contrast, motion coherence, and translational speed were high. Under explicit tracking instruction, the eyes aligned with the focus of expansion more closely than without instruction. Our results highlight the sensitivity of intuitive eye movements as indicators of visual motion processing in dynamic contexts.

## Introduction

Many daily functions, from catching prey to walking or driving to work, involve moving through our dynamic visual environment. To adequately control self-motion, we need to know not only whether we are moving, but also where and how fast. These aspects of self-motion are informed by multiple sensory cues, one of which is optic flow, the visual motion pattern projected onto our retina when we move through the visual environment (Gibson, 1950; Lappe, Bremmer, & van den Berg, 1999; Vaina, Beardsley & Rushton, 2004). Optic flow expands or radiates outward when we move forward, and contracts or radiates inward when we move backward. The singular point of convergence or radiation is termed the focus of expansion (FOE) and often indicates heading direction.

In humans and other animals, optic flow is critically important in guiding locomotion (Gibson, 1958; Warren, Kay, Zosh, Ducho & Sahuc, 2001), posture maintenance (e.g., Bardy, Warren, & Kay, 1996; 1999), and navigation such as steering (e.g., Li & Niehorster, 2014). Other animals also adjust their locomotive or flight behavior based on optic flow (insects: Srinivasan & Zhang, 2000; birds: Bhagavatula, Claudianos, Ibbotson, & Srinivasan, 2011; Dakin, Fellows, & Altshuler, 2016). Accordingly, humans and macaque monkeys are able to reliably discriminate FOE position changes as small as 1 degree (°) of visual angle (Britten & Wezel, 2002; Warren & Hannon, 1988). The ability to perceive and discriminate the FOE position depends on where observers look (Warren & Kurtz, 1992; Crowell & Banks, 1993; Gu, Fetsch, Adeyomo, DeAngelis, & Angelaki, 2010). These studies tested heading direction discrimination thresholds while observers fixated at different eccentricities relative to the FOE. They found that performance scaled as a function of proximity to the fovea, with the best performance when the FOE was near the fovea.

Viewing an optic flow stimulus with a stationary and eccentric FOE generates eye movements toward the FOE (macaque monkeys: Lappe, Pekel, & Hoffmann, 1998, 1999; Angelaki & Hess, 2005; humans: Niemann, Lappe, Büscher & Hoffmann, 1999). These studies showed that observers make a combination of saccades directed to the FOE, as well as slow-tracking reflexive eye movements following retinal motion near the fovea. Recently, Knöll, Pillow, and Huk (2018) found that different primate species – one human, two macaques, and one marmoset monkey – intuitively tracked a dynamically changing FOE with their eyes despite minimal training or instruction. In conjunction with evidence showing that eye movements can be more sensitive than motion perception (Tavassoli & Ringach, 2010; Spering & Carrasco, 2015), these recent studies emphasize the value of eye movements as sensitive indicators of the processing of visual motion features.

Notwithstanding the known sensitivity of eye movements and the finding that the eyes naturally move to the FOE across species (Knöll et al., 2018; Spering & Chow, 2018), we do not yet know how often or consistently observers lock their gaze onto the FOE, or how accurate these intuitive eye movements are. A better characterization of eye movements in such a naturalistic context could lead to a better understanding of how eye movements might serve heading and locomotion. Moreover, intuitive eye movements are potentially a powerful tool for clinical testing and diagnosis, where patients can often not be explicitly instructed. Understanding how these eye movements respond to different tasks and visual constraints could be a stepping stone toward the development of eye-movement based tools to investigate motion sensitivity. Here we characterize eye movement responses toward an optic flow stimulus with an unpredictably moving FOE.

First, we will investigate whether FOE is tracked intuitively by a large sample of human observers with a simple instruction to free-view the stimulus. If so, this will confirm previous observations in a small sample (Knöll et al., 2018). We will test this by quantifying the occurrence and variability of intuitive FOE-tracking behavior through the analysis of the overall alignment of the eye and FOE position changes, smooth tracking (optokinetic reflex or smooth pursuit), and fast tracking (saccades).

Second, we will investigate how eye movements change when observers receive explicit instruction to track the FOE versus when they are uninstructed. Whereas psychophysical testing in the laboratory usually involves explicit task instructions combined with feedback to promote task compliance, natural stimulus encounters do not involve explicit instruction. Moreover, the ability to understand and follow instructions depends on an observer’s cognitive state. Here we will quantify the potential improvement of FOE-tracking measures by explicit instruction.

Third, we will investigate how FOE tracking changes when we manipulate motion signal strength by altering stimulus features such as contrast, coherence, and speed. These manipulations mimic daily environments where we regularly experience visual motion in a range of signal strengths, such as when driving through fog or moving slowly in heavy traffic. Previous studies have demonstrated how these features affect visual motion processing in trained observers under explicit instruction. For instance, observers’ accuracy in discriminating between optic flow direction increased with increasing stimulus coherence (~25% coherence required for a brief stimulus of 100 ms, Burr & Santaro, 2001). Increasing luminance contrast did not improve discrimination performance beyond a certain level (e.g., 3%, Allen, Hutchinson, Ledgeway, & Gayle, 2010; 15%, Edwards, Badcock, Nishida, 1996), suggesting early saturation of contrast in motion processing. When instructed to track the FOE with their eyes, observers showed improved spatial and temporal alignment when speed increased from 1 m/s to 4 m/s and from 4 m/s to 16 m/s (Cornelissen & van den Dobbelsteen, 1999). Analyzing the impact of motion signal strength on intuitive FOE tracking will address whether intuitive eye movements reveal similar characteristics of motion processing previously established via instructed tasks.

## Method

In two experiments, we characterized intuitive tracking performance. In Experiment 1, we asked observers to view a high-contrast, high-coherence optic flow stimulus under free-viewing. In Experiment 2, we manipulated stimulus signal strength (low vs. high coherence, contrast, and speed) and instruction (free viewing vs. explicit instruction to the track the FOE).

### Observers

We tested 43 adults (age mean = 25.7, +/- 5.0 years; 22 female); *n* = 19 participated in Experiment 1 and *n* = 24 in Experiment 2. All observers provided data in the high-contrast, high-coherence condition, while each subset of observers provided additional data in our stimulus and instruction manipulations (*n* ≥ 19 for each comparison). These sample sizes are considerably larger than sample sizes used in similar studies in the literature (e.g., Knöll et al., 2018: *n* = 1; Niemann et al., 1999: *n* = 4). Due to problems with the eye-tracking setup, three observers did not complete all conditions after instruction, resulting in different *n* when comparing the effect of motion signal strength (*n* = 22 for motion coherence; *n* = 21 for dot contrast; *n* = 22 for translational speed). All observers had normal or corrected-to-normal visual acuity (20/20), tested using a Snellen visual acuity chart. Observers received CAD 12 per hour as remuneration for their participation. The experiment protocol adhered to the Declaration of Helsinki and was approved by the University of British Columbia Behavioral Research Ethics Board. Before participation, observers provided written informed consent.

### Visual Display and Apparatus

In both experiments, observers were seated 55 cm away from a 39 cm × 29 cm CRT monitor (ViewSonic G255f; resolution 1280 × 1024 pixel; refresh rate 85 Hz) covering 39.0° × 29.5° of the visual field. Each observer’s head was stabilized using a combined chin- and forehead-rest. Visual stimuli were generated by a PC with an NVIDIA GeForce GRX 970 graphics card, using Matlab R2018b (MathWorks), Psychtoolbox 3.0.12 (Brainard, 1997; Kleiner et al., 2007; Pelli, 1997), and PLDAPS toolbox version 4 (Eastman & Huk, 2012).

### Stimuli

Our stimulus was similar to that used in Knöll and colleagues (2018) and was comprised of a cloud of 295 dots at a density of 0.4 dots/deg^2^ covering 31.2° × 23.6° of the visual field. Each dot was 0.13° × 0.13° in size and lasted for an unlimited lifetime (Exp. 1) or a short lifetime of 4 frames (47 ms; Exp. 2). The background had a luminance of 50 cd/m^2^; in Experiment 1, half of the dots were 33% brighter, and half of the dots were 33% darker than the background (Michelson contrast). In Experiment 2, dot Michelson contrast was set to one of five contrast levels (3.6, 8.1, 18.1, 40.3, 90%) when contrast was manipulated, and to 90% contrast when either motion coherence or speed was manipulated.

Each dot was categorized as a signal dot or a noise dot to achieve a desired global motion coherence level. In Experiment 1, the motion coherence level was 100%; in Experiment 2, motion coherence was set to one of five coherence levels (6.25, 12.5, 25, 50, 100%) when coherence was manipulated, and to 100% coherence when contrast or speed was manipulated. Noise dots moved randomly, whereas the signal dot location was updated based on the 3D velocity of the dot cloud. The 3D velocity of the dot cloud depended on the simulated translational speed: dots moved more quickly when the simulated translational speed was higher, and vice versa. In Experiment 1, translational speed was constant at 2 m/s; in Experiment 2, translational speed was set to one of five levels (0.75, 1.5, 2, 3, 6 m/s) when speed was manipulated, and to 2 m/s when coherence or contrast were manipulated. The in-depth motion direction of the dot cloud alternated randomly, resulting in expanding or contracting optic flow, with an average switch time of 3.6 s. The 3D velocity of the dot cloud also depended on the location of the FOE on the screen. In general, dots further away from the FOE moved more quickly (maximum velocity 78°/s), and dots at the FOE were stationary.

The FOE location shifted continuously in the horizontal and vertical dimensions in a random-walk fashion. The movement shift was small, yielding a location variability of *SD* = 2.7° within a given trial, in Experiment 1, and larger (*SD* = 4.5°) in Experiment 2 to increase the demand of active tracking. The statistical dynamics of the FOE movement were kept constant across manipulations of motion signal strength listed above, i.e. even when the dot speed was manipulated, the speed of the FOE they encoded did not alter between these trials.

### Procedure and Design

In Experiment 1, observers completed one free-viewing block of 3-3.5 minutes accumulated viewing time with alternating optic flow direction. In Experiment 2, observers always first completed three blocks of free-viewing, one for each motion signal strength manipulation (contrast, coherence, speed, in randomized order), followed by three blocks (one block each for contrast, coherence, speed) in which they were asked to track the FOE with their eyes. Each block consisted of five trials of 30 s duration each. Random seeds were used between blocks and observers; the same seed was used within each block such that all trials within a block shared the same FOE position trajectory.

In both experiments, stimulus presentation was triggered when the observer looked at the screen and paused if they looked near the edge or outside the screen or blinked for more than 250 ms. Observers did not receive feedback regarding tracking performance. To ensure that all observers had the same level of exposure to the optic flow stimulus before the experiment started, they first viewed an exposure trial (30 s) with an obvious optic flow stimulus (high motion coherence, high dot contrast, medium speed) with free-viewing instruction.

### Eye Movement Preprocessing and Analysis

Eye position data were recorded at 1000 Hz using a video-based eye tracker (Eyelink 1000; SR Research Ltd., Ottawa, Canada) and processed offline using Matlab. Eye position trajectory (down-sampled to 850 Hz) was compared with the FOE position trajectory (up-sampled to 850 Hz). Eye position data were filtered using a Butterworth filter (low pass, second-order) with a cutoff frequency of 15 Hz for position and 30 Hz for velocity. Blinks and eye position data 50 ms before and after blinks were removed from all analyses (5.2% of position data across observers).

**Table 1** summarizes measures used to describe FOE-tracking behavior. To assess the overall alignment between the FOE and eye position changes across time, we conducted cross-correlation analyses between each observer’s eye position and FOE position across all samples obtained in a trial, where correlation coefficients were generated after introducing a variable time lag between the eye and FOE position. Based on results presented in Knöll et al. (2018), we limited the correlation analysis by the following constraints: the eye had to lag behind the FOE (not be ahead of it), and the maximum lag was 10 s. For each observer, we then used the highest yielded correlation coefficient as an indicator of overall alignment; higher correlation coefficient equals better alignment. To quantify temporal alignment, we used the time lag that yielded the highest correlation coefficient; the shorter the time lag, the faster the eyes caught up with FOE position changes. To quantify spatial alignment at zero lag, we used the average Euclidean distance between the eye and FOE position across time frames; the smaller the position error, the more closely the eyes hovered around the FOE.

**Table 1.**
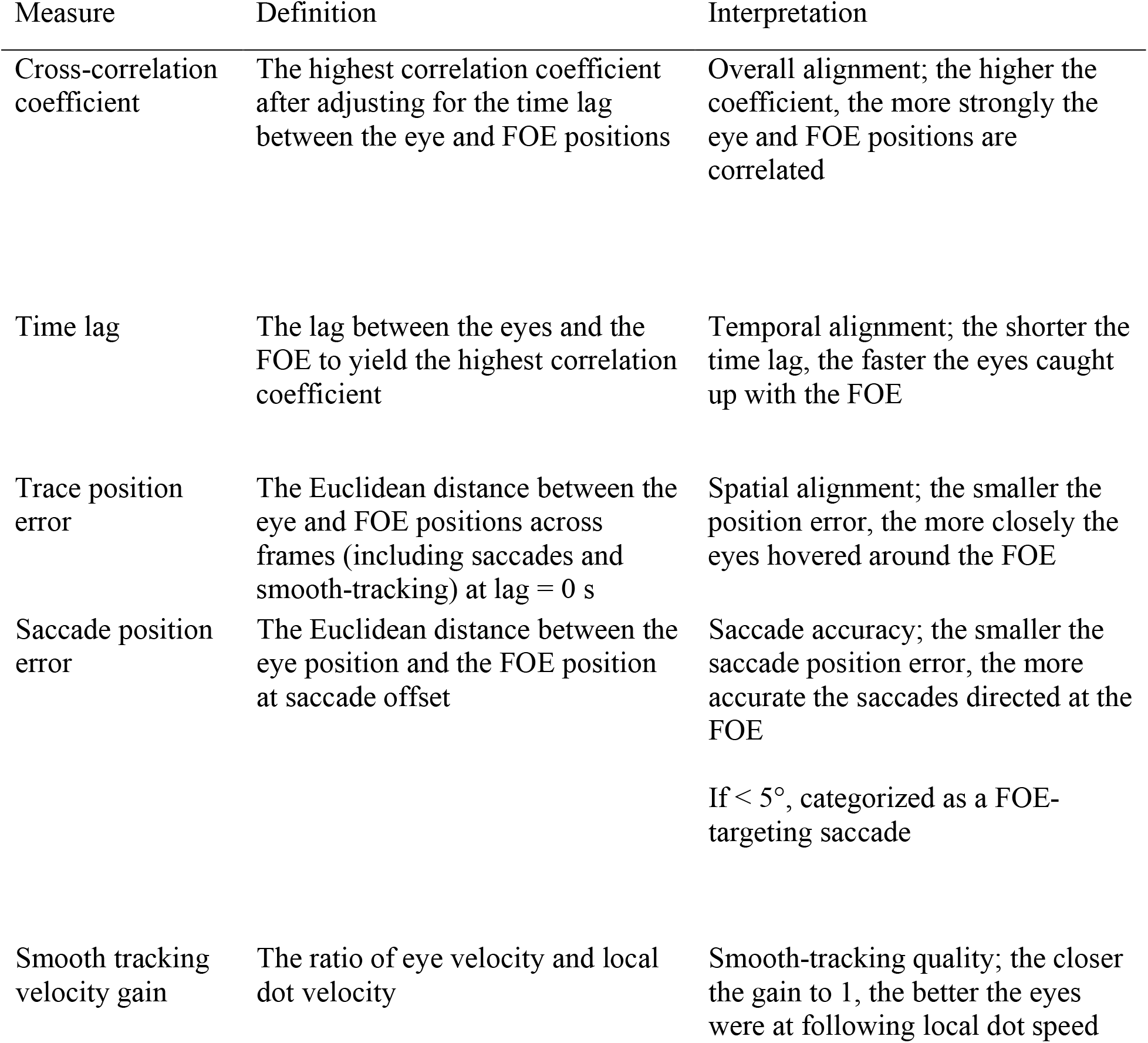
A summary of analysis measures to describe the temporal and spatial accuracy of eye movements relative to the FOE

| Measure | Definition | Interpretation |
| --- | --- | --- |
| Cross-correlation coefficient | The highest correlation coefficient after adjusting for the time lag between the eye and FOE positions | Overall alignment; the higher the coefficient, the more strongly the eye and FOE positions are correlated |
| Time lag | The lag between the eyes and the FOE to yield the highest correlation coefficient | Temporal alignment; the shorter the time lag, the faster the eyes caught up with the FOE |
| Trace position error | The Euclidean distance between the eye and FOE positions across frames (including saccades and smooth-tracking) at lag = 0 s | Spatial alignment; the smaller the position error, the more closely the eyes hovered around the FOE |
| Saccade position error | The Euclidean distance between the eye position and the FOE position at saccade offset | Saccade accuracy; the smaller the saccade position error, the more accurate the saccades directed at the FOE<br>If $< 5^\circ$, categorized as a FOE-targeting saccade |
| Smooth tracking velocity gain | The ratio of eye velocity and local dot velocity | Smooth-tracking quality; the closer the gain to 1, the better the eyes were at following local dot speed |

The random FOE-shifts triggered a combination of different types of eye movements, saccades and smooth tracking. Saccadic eye movements were detected when the eye velocity (the digitally differentiated eye position) exceeded a fixed velocity criterion (30 deg/s) for five consecutive frames. Acceleration minima/maxima determined saccade onsets/offsets. In addition to saccade duration, amplitude, and direction, we quantified the relationship between the saccade and the FOE by position error as a measure of spatial accuracy measure. We defined saccade position error as the Euclidean distance between eye position at saccade offset and FOE position at the time of saccade offset; lower position error equals higher accuracy. The quality of smooth-tracking eye movements during inter-saccade intervals was characterized by analyzing their peak and median eye velocity, and their dot velocity gain (ratio of eye velocity vs. local dot velocity near the fovea); higher velocity gain equals better quality. Short inter-saccade intervals (< 50 ms) or extremely long intervals (3 *SD* or more than the mean inter-saccadic interval duration for each observer) were removed from the analysis (2.3% of the inter-saccade intervals across observers).

When observers did not track, random eye movements could still land onto the FOE by chance. To differentiate tracking from tracking that occurred by chance (tracking-at-chance), we computed baseline alignment and eye movement measures that indicated the expected values for tracking-at-chance. This analysis was achieved by comparing observers’ eye positions with secondary FOE positions that were not used to generate the flow pattern, despite sharing the same statistical dynamics as the FOE shown, thus indicating performance when tracking was at chance. From a total of 474 comparisons across 30 s each, this analysis yielded a distribution of baseline measures.

### Statistical Analysis

For each analysis, we first tested whether the relationship between variables was different between Experiments 1 and 2; if no differences were observed, data were collapsed across experiments. We also assessed for differences between responses to horizontal and vertical stimulus dimensions, and limit reporting results to the horizontal dimension when no differences were observed. To quantify the occurrence and variance of intuitive FOE tracking, we report descriptive statistics, such as median (*Mdn*) and standard deviation (*SD*), of alignment measures (cross-correlation coefficients, time lag, trace position error) as well as eye movement measures (saccade position accuracy and smooth-tracking quality) across observers in Experiments 1 and 2. To compare intuitive FOE tracking to chance, the *Mdn* and *SD* of the baseline measures of tracking-at-chance were reported for reference. Furthermore, observers were categorized as intuitive trackers if their cross-correlation coefficients under free-viewing were at least 2 *SD* above the group *Mdn*. To investigate whether tracking depended on motion signal strength (high vs. low dot contrast, motion coherence, global translational speed) and instruction (free viewing vs. tracking), we compared saccades and smooth tracking between conditions using the Wilcoxon signed-ranks test and Friedman test, respectively. When a factor with more than two levels was found significant, a post-hoc Wilcoxon signed-ranks test adjusted for multiple comparisons by false-discovery rate was performed. Additional analyses that might be of interest to readers, such as the effect of time on task and optic flow directions (expansion vs. contraction) in Experiment 1, as well as the interaction between signal strength and instruction manipulations in Experiment 2, were reported in **Supplementary Materials**. All statistical analyses were conducted in R (R Core Team, 2017). Non-parametric statistical tests were used because data were not normally distributed based on visual inspection. All result figures were produced using R package ggplot2 (Wickham, 2009).

## Results

### Human observers intuitively track the FOE with their eyes

A comparison of eye position to FOE position shows a good temporal and spatial correspondence between the eye and the FOE when observers viewed an optic flow stimulus of high motion-signal strength. **Figure 1** shows eye position trajectories from two individual observers in one trial, whereas **Figure 2** shows the tracking performance of all 43 observers.

**Figure 1.**
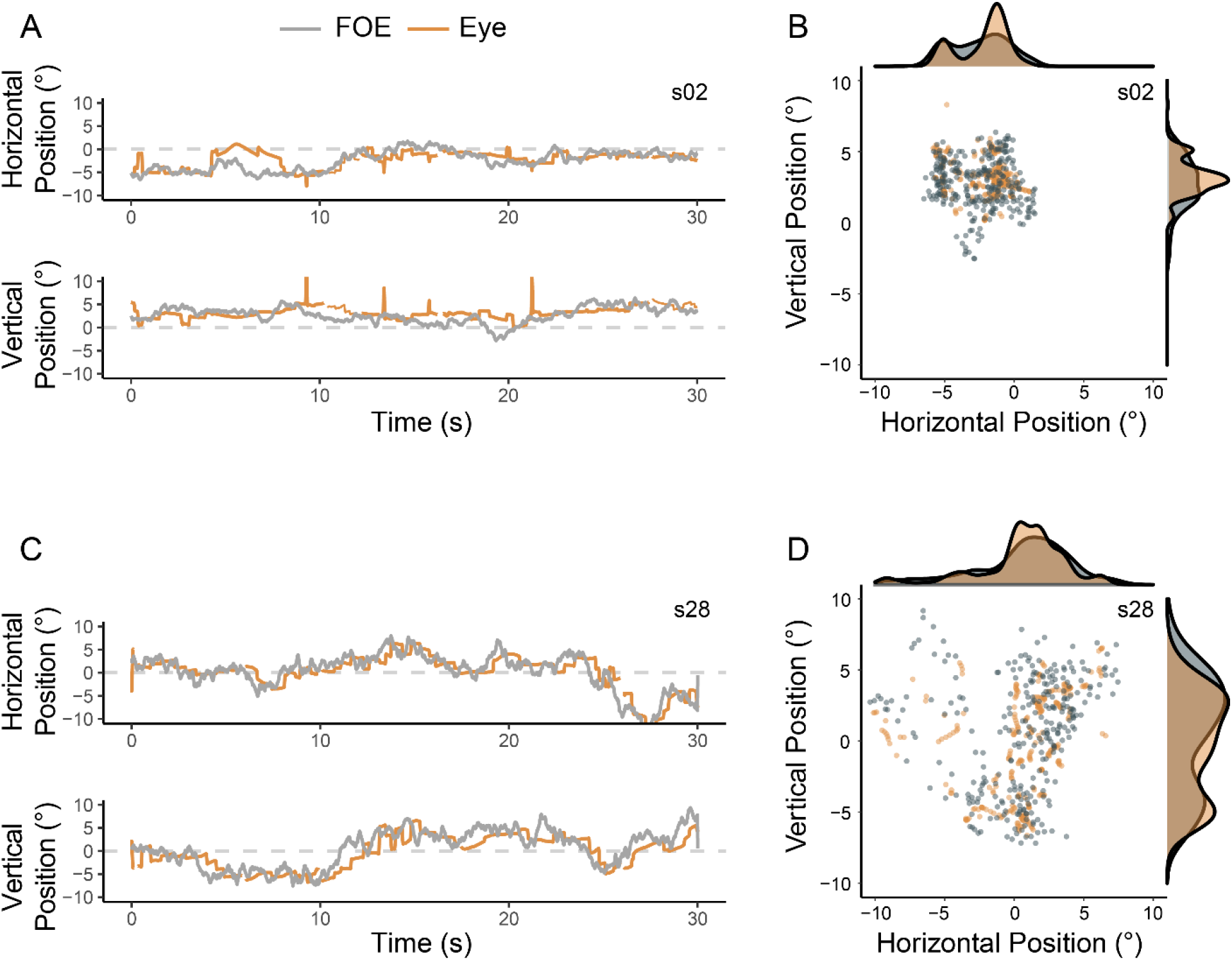
Horizontal and vertical position trajectories of the FOE (grey line) and the eyes (orange line) of two representative observers (A,B) and (C,D) showing a good temporal and spatial correspondence between eyes and FOE. Panels on the right show comparisons of the spatial distribution of eye positions and FOE in the same trials depicted on the left. Each data point represents the average in a 100-ms interval, yielding a total of 300 data points plotted for each observer across a period of 30 s. Probability density plots of horizontal and vertical positions were plotted respectively above and to the right of panel B, D.

**Figure 2.**
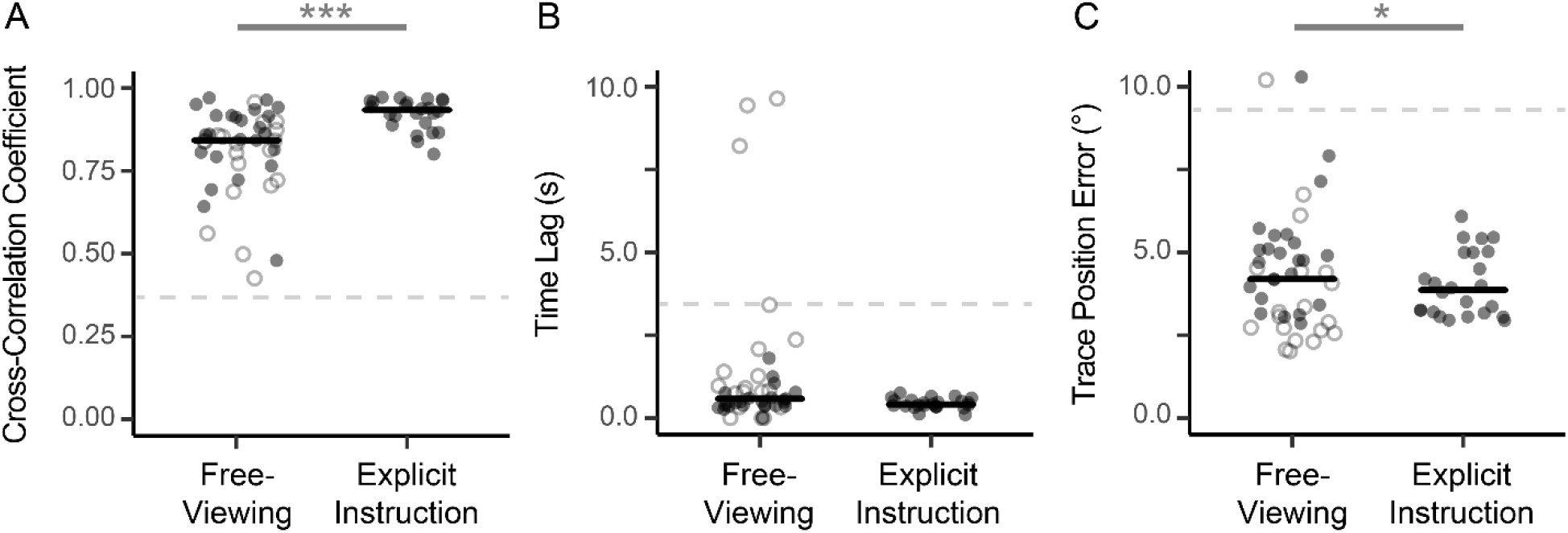
A quantification of FOE-tracking behavior under free-viewing (*n* = 43) and explicit instruction (*n* = 24). Data from Experiment 1 and Experiment 2 are plotted in open and solid dots, respectively. Positions along the horizontal axis were jittered to reduce overlapping in data points. The horizontal black line shows the median across observers and experiments. The grey dashed line indicates the median baseline measures of tracking-at-chance (cross-correlation coefficient: 0.34; time lag: 3.27 s; trace position error: 8.1°). Asterisks indicate *p*-values in pairwise comparisons based on the subset of observers completing both conditions in Experiment 2 (*: *p* < .05, ***: *p* < .001).

In terms of temporal alignment between eye and FOE, the eyes tracked the FOE with a median cross-correlation coefficient of 0.84 (*SD* = 0.13, **Fig. 2A**) at a median time lag of 581 ms (*SD* = 2.26 s, **Fig. 2B**) across observers. This temporal alignment is reflected in the example observers’ eye position trajectories (**Fig. 1A, 1C**), indicated by small systematic rightward shifts (s02: 500 ms, s28: 336 ms) along the time axis for the eye position trace. The shape of the position traces – how the position changed as a function of time – was highly similar across both eye and FOE positions, yielding a high (closer to one) cross-correlation coefficient (s02: 0.83, s28: 0.94). Both observers had a cross-correlation coefficient of at least 2 *SD* above the simulated baseline cross-correlation coefficient (*Mdn* = 0.34, *SD* = 0.18). If we consider cross-correlation coefficients of ≥ 0.7 (2 *SD* above median baseline measure) as indicators of good tracking, 36 out of 43 observers (84% of our sample) showed intuitive FOE-tracking when given free-viewing instruction.

In terms of spatial alignment, the median trace position error across all observers was 4.2° (*SD* = 1.9°; **Fig. 2C**). **Figures 1B** and **1D** reveal an overlap between 2D eye position and FOE position in the two example observers, indicated by density plots showing the frequency distribution of horizontal and vertical eye and FOE positions, respectively. The spread of the eye horizontal position distributions matched the respective FOE horizontal position distributions well with >60% of overlap for the two selected observers (s02: 64%, s28: 76%), even though a simple permutation test revealed statistical differences between the distribution shapes for one of the two observers (s02: *p* < .001, s28: *p* = .058).

To investigate how observers achieved accurate FOE tracking, we analyzed saccades in more detail. Across experiments, observers made a median of 1.3 saccades/s (*SD* = 0.46 saccades/s) with an amplitude of *Mdn* = 2.4° (*SD* = 0.9°) and a duration of *Mdn* = 98 ms (*SD* = 20.6 ms). Across observers, the majority (71%) of these saccades (equivalent to 0.9 saccades/s) landed within 5° of the FOE with a median position error of 3.4° (*SD* = 1.9°). For smooth-tracking eye movements, observers’ eyes moved slowly at an average velocity of 2.1°/s (*SD* = 0.8°/s) and a dot velocity gain of 0.29 (*SD* = 0.12). Both saccade accuracy and smooth-tracking quality were higher than baseline performance of tracking-at-chance (baseline saccade position error: *Mdn* = 7.7°, *SD* = 5.0°; baseline smooth-tracking velocity gain: *Mdn* = 0.19, *SD* = 0.47).

These results show that most observers intuitively tracked the FOE with high accuracy, suggested by the good temporal and spatial correspondence between the FOE and the eyes. Observers did so by directing their saccades at the FOE, indicated by a large proportion of saccades landing close to the FOE. This tracking performance was extracted from only 30 s of viewing time (see **Supplementary Analysis and Figure S1**). Tracking performance was comparable across optic flow directions (see **Supplementary Analysis and Figure S2**).

### Explicit instructions improve FOE-eye alignment and saccade accuracy

Explicit tracking instruction improved the overall alignment between the eyes and the FOE across all observers who participated in both instruction conditions (**Fig. 2**; solid dots, *n* = 24). The overall FOE-eye alignment improved by 8.1% with explicit instruction, indicated by an increase in cross-correlation coefficient (instruction: *Mdn* = 0.93, *SD* = 0.05; free-viewing: *Mdn* = 0.86, *SD* = 0.12; *Z* = 34, *p* < .001, *r* = .68, **Fig. 2A**). We also observed a 15% decrease in trace position error in trials with explicit instruction (*Mdn* = 3.9°, *SD* = 1.0°) vs. with free-viewing instruction (*Mdn* = 4.6°, *SD* = 1.7°; *Z* = 233, *p* = .016, *r* = .48, **Fig. 2C**), and a 11% reduction in time lag, from 459 ms (*SD* = 367 ms) with free-viewing instruction to 409 ms (*SD* = 158 ms) with explicit instruction, which was not significant (*Z* = 211, *p* = .08, *r* = .36, **Fig. 2B**). These findings suggest that explicit instruction improved the overall mapping between the eyes and the FOE in time and in space, but not necessarily how fast the eyes catch up with the FOE.

Explicit instruction improved saccade accuracy. Under explicit instruction, 77% (*SD* = 15%) of all saccades were FOE-targeting saccades with a median position error of 3.4° (*SD* = 0.8°). During free viewing, observers made significantly fewer targeting saccades (*Mdn* = 62%, *SD* = 21%; *Z* = 64, *p* = .013, *r* = .50) and saccade position error was significantly higher (*Mdn* = 4.1°, *SD* = 1.6, *Z* = 239, *p* = .010, *r* = .52), as shown in **Figure 3A**. Saccade rate did not differ between instruction (*Mdn* = 1.5 saccades/s, *SD* = 0.4 saccades/s) and free viewing (*Mdn* = 1.4 saccades/s, *SD* = 0.5; *Z* = 114, *p* = .31, *r* = .21).

**Figure 3.**
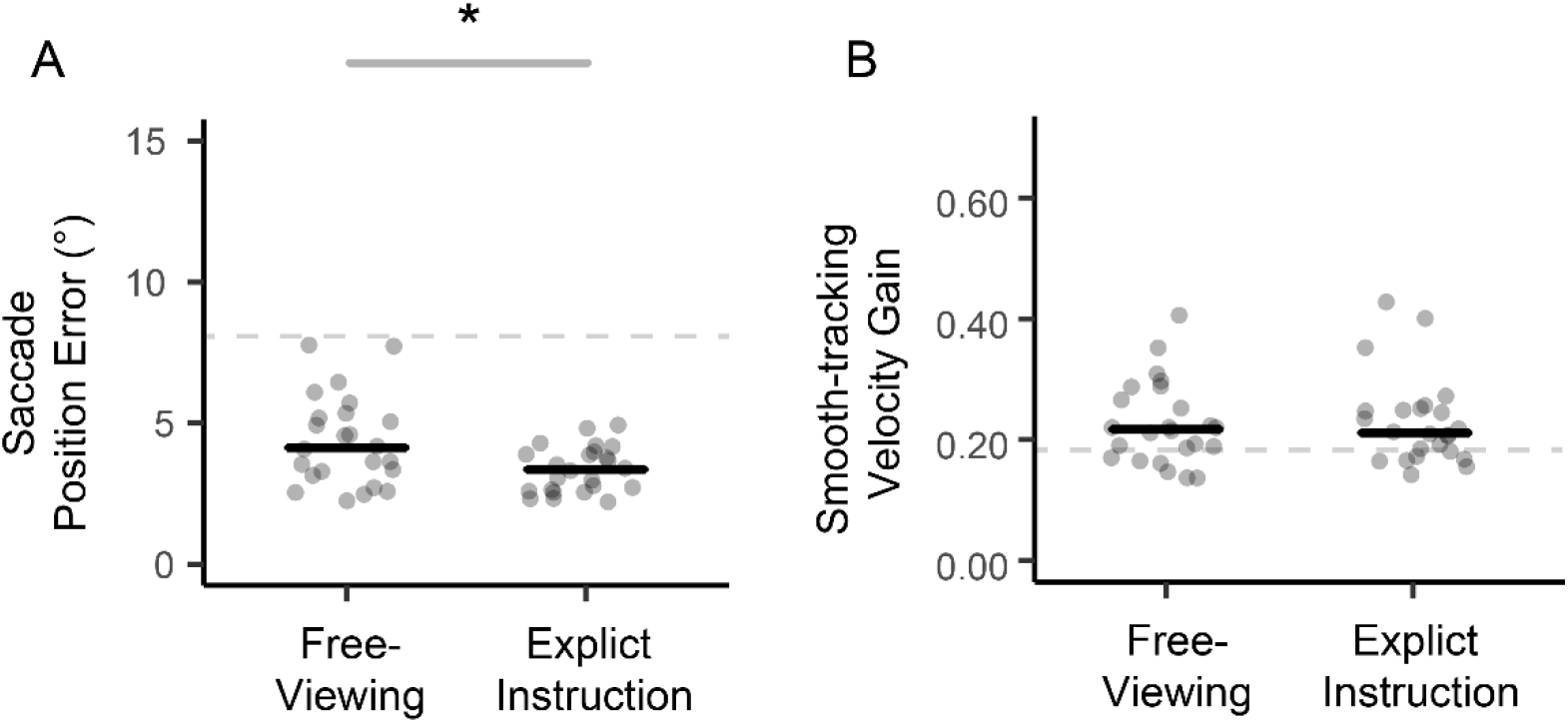
The effect of explicit instruction on saccade position error (A) and smooth-tracking velocity gain (B) in Experiment 2. Individual data points were plotted in gray shapes with the positions jittered along the horizontal axis to reduce overlap. The horizontal black line shows the median across observers. The grey dashed line indicates the median baseline measures of tracking-at-chance (saccade position error: 7.7°; smooth-tracking velocity gain: 0.19). Asterisks indicate *p*-values in pairwise post-hoc comparisons (*: *p* < .05).

By contrast, smooth-tracking performance was similar across free-viewing and explicit instruction conditions as shown in **Figure 3B**. Under explicit instruction, observers’ eyes moved at a median velocity of 1.9°/s (*SD* = 0.6°/s) and at a dot velocity gain of 0.21 (*SD* = 0.07). These measures are not statistically different from measures obtained during free viewing (velocity: *Mdn* = 2.2°/s, *SD* = 0.7°/s, *Z* = 203, *p* = .14, *r* = .31; gain: *Mdn* = 0.21, *SD* = 0.07, *Z* = 170, *p* = .58, *r* = .12).

Taken together, these results show that explicit instruction improved the correspondence between the eyes and the FOE via improvements in saccade accuracy, but not improvements in smooth tracking. **Supplementary Analysis and Figure S3** further show that this is the case at most signal strength levels.

### Intuitive eye movements scale with stimulus signal strength

We next investigated effects of motion signal strength on the accuracy and quality of intuitive eye movements under free-viewing instruction. As shown in **Figure 4A-C**, saccade position error was generally reduced when motion signal strength increased, confirmed by a significant main effect of motion signal strength in all three stimulus manipulations (coherence: *χ^2^*(4) = 16.7, *p* = .002, *W_Kendall_* = .19; contrast: *χ^2^*(4) = 12.1, *p* = .017, *W_Kendall_* = .14; speed: *χ^2^*(4) = 42.9, *p* < .001, *W_Kendall_* = .49). Similarly, smooth-tracking velocity gain (**Fig. 4D-F**) was modulated by motion signal strength in all stimulus manipulations (coherence: *χ^2^*(4) = 48.9, *p* <.001, *W_Kendall_* = .56; contrast: *χ^2^*(4) = 40.4, *p* < .001, *W_Kendall_* = .48; speed: *χ^2^*(4) = 57.9, *p* <.001, *W_Kendall_* = .66).

**Figure 4.**
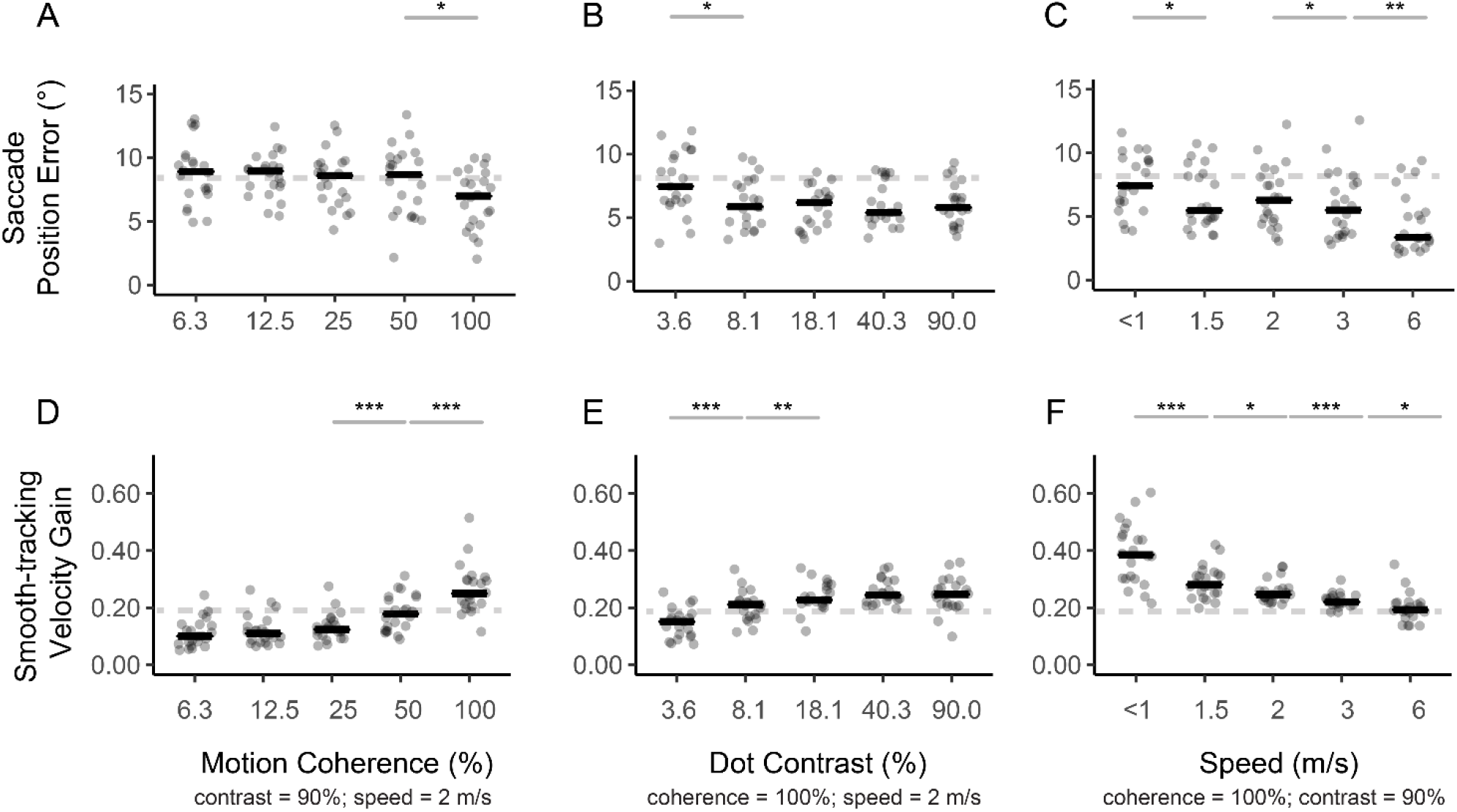
The influence of motion signal strength (horizontal axis) on saccade position error (top row) and smooth-tracking velocity gain (bottom row) in Experiment 2 under free-viewing, where motion coherence (A,D), dot contrast (B,E), and global translational speed (C,F) were manipulated respectively. Individual data points were plotted in gray shapes with the positions jittered along the horizontal axis to reduce overlap. The horizontal black line shows the median across observers. The grey dashed line indicates the median baseline measures of tracking-at-chance (saccade position error: 7.7°; smooth-tracking velocity gain: 0.19). Asterisks indicate *p*-values in pairwise post-hoc comparisons (*:*p* < .05, **: *p* < .01, ***: *p* < .001).

A closer examination showed that saccade position error and velocity gain did not change parametrically at each step of motion signal strength. For motion coherence (**Fig. 4A**), both measures showed the largest change at the highest level of motion coherence, reflected in a significant pairwise comparison when motion coherence increased from 50 to 100%. For dot contrast (**Fig. 4B**), both measures showed the largest change at the lowest level of dot contrast, when dot contrast increased from 3.6% to 8.1%. For translational speed (**Fig. 4C**), saccade accuracy increased, and smooth-tracking quality decreased significantly at each increment of speed.

These results show that saccade accuracy and smooth-tracking quality under free-viewing were sensitive to motion signal strength. In general, the higher the motion signal strength, the lower the saccade position error, as well as the higher the velocity gain. Our results also highlight differences in stimulus manipulations. Among the chosen stimulus values, changes in eye movement measures were most apparent when motion coherence was high and when dot contrast was low, whereas translational speed induced changes in eye movement measures across all speed levels. In **Supplementary Analysis and Figure S3**, we show that the effects of motion signal strength were similar after explicit instruction, with changes in eye movement measures being more pronounced compared to free-viewing. These results suggest that eye movements are sensitive indicators of motion signal characteristics and might reflect more general motion processing mechanisms.

## Discussion

In this study, we characterized eye movement responses triggered by optic flow in a larger group of human observers and report the following key findings. First, over 80% of observers intuitively tracked a dynamically moving FOE. This tracking was achieved by a combination of saccades directed to the FOE and smooth-tracking eye movements in response to local dot motion near the fovea. Second, explicit instruction improved the overall correspondence between the eye and the FOE via improvements in saccade accuracy, but not the quality of smooth-tracking eye movements. Third, intuitive eye movement measures depended on motion signal strength. Taken together, these findings show that intuitive eye movements triggered by optic flow allow us to probe how the visual system processes dynamic visual scenes relevant to self-motion. In the following paragraphs, we will discuss the consistency of this intuitive alignment between the eyes and the FOE with previous studies, how this alignment is achieved by using a combination of different types of eye movements, and its potential applications as sensitive indicators of performance in various dynamic tasks.

### The spatial and temporal alignment between eyes and FOE are consistent across studies

Self-motion, object motion, and eye movements interact to create a dynamic visual scene, including the shifting of the FOE. In our study, observers used a combination of saccades and smooth tracking to align the fovea with the FOE with a median eye position error of 4.2°, congruent with previous research. For example, real-world eye-tracking studies showed that human observers looked at the lead car or the center of the lane while driving (Mourant, Rockwell & Rackoff, 1969; Rockwell, 1972), with 90% of the observed fixations within 4° of the FOE. Using a saccade task, Hooge, Beintema, and van den Berg (1999) showed that observers’ saccade endpoint scattered within 4° of the FOE in diameter. When viewing an optic flow stimulus with a FOE shifting sinusoidally in the horizontal dimension, observers locked their gaze on average within 4.6° of the FOE for about 70% of the time (Shirai & Imura, 2016). The similarity between spatial performance across studies suggests that our normative performance can be generalizable across stimuli (real-world visual scene vs. laboratory-generated optic flow), tasks (instructed saccade task vs. free-viewing task), and FOE movement types (predictable one-dimensional vs. unpredictable two-dimensional). Intuitive performance is highly relevant in understanding real-world behavior across settings.

In the temporal domain, our analysis showed that under free-viewing instruction the eyes lagged behind the FOE by a median of 581 s (considering the entire sample of naïve observers) or 561 ms (considering only the sample who tracked intuitively), both of which are longer than similar measures reported in previous research. However, when considering a subset of data based on explicit instruction, the average time lag dropped to 436 ms, comparable to previous studies. For example, Knöll and colleagues (2018) reported in one human observer that the eyes track the FOE with a lag of 300 ms. Cornelissen and van den Dobbelsteen (1999) showed that a small sample of instructed observers tracked the FOE with a time lag of 300-450 ms. When an optic flow stimulus was briefly shown for ~300 ms, observers can reliably place the cursor near the FOE (te Pas, Kappers, & Koenderink, 1998) or discriminate between stimuli of different heading directions (Crowell et al., 1990). Accounting for saccade dead time (when saccades cannot be altered), Hooge et al. (1999) estimated that the processing time of heading direction is 430 ms based on saccade latency and saccade landing error. Overall these studies suggest that heading direction can be extracted in under 450 ms, consistent with what we found in trials in which observers were instructed to track. In sum, spatial and temporal alignment between the eyes and the FOE shows remarkable similarities with previous reports, suggesting that intuitive tracking can capture performance across settings.

### Saccades and pursuit jointly support intuitive tracking

To achieve this consistent and intuitive tracking of the FOE, observers in our study exhibited a combination of saccades and smooth-tracking eye movements similar to previous reports (Lappe et al., 1998; Angelaki & Hess, 2005; Knöll et al., 2018; Lappi, Pekkanen, Rinkkala, Tuhkanen, Tuononen, & Virtanen, 2020). This combined behavior is aligned with the notion that these eye movements are complementary for visual tracking of unpredictable targets (Orban de Xivry & Lefèvre, 2007). Whereas the overall saccade rate in our study was lower than the saccade rate in other studies using free-viewing instructions of natural scenes (2.9 saccades/s, Otero-Millan, Troncoso, Macknik, Serrano-Pedraza, & Martinez-Conde, 2008), saccade endpoints were aligned with the FOE, yielding an observed percentage of FOE-targeting saccades similar to a previous study (Knöll et al., 2018). When observers were not making saccades, they exhibited smooth-tracking eye movements at low velocity. The gain of smooth-tracking (0.35 with unlimited dot lifetime in Exp 1; 0.21 with a short dot lifetime of 47 ms in Exp 2) was lower than the gain (approx. 0.6) achieved in other studies for passive viewing of optic flow (Niemann et al., 1999, dot lifetime not reported). The discrepancy might be partly attributed to the short dot lifetime adopted in Experiment 2, as evident by the difference in gain across experiments. However, even with unlimited lifetime, the gain in Experiment 1 was still low. The low velocity-gain in our study suggests that observers did not strategize to track local dots. Otherwise, their velocity gain would have easily reached unity, as demonstrated by Niemann et al. (1999), after explicit instruction to follow local dot motion. The results obtained for combined saccade and pursuit tracking suggest that our saccadic eye movement system is highly responsive to global features of a complex visual stimulus, like the FOE. Whereas the pursuit system is optimized for tracking moving objects, it merely plays a supplementary role in FOE-tracking. Extracting the FOE is more challenging than tracking a small, moving dot with smooth pursuit, or than determining the direction of a homogenously moving field with optokinetic nystagmus. Furthermore, the random shifts of FOE movement also mean that tracking cannot be perfect, because observers cannot predict when and where the next shift would be. These differences might explain why even under explicit instruction, the eyes still deviated from the FOE position by 4° and tracked at a gain of only 0.21.

### Intuitive tracking is sensitive to stimulus features

Our study further shows that intuitive eye movements to the FOE are sensitive to all three stimulus features manipulated, albeit being affected to a different extent. These results resemble known characteristics of visual motion processing established in previous literature from instructed observers. A lower motion coherence indicates that a smaller proportion of dots is moving in a direction corresponding with the FOE. A lower translational speed indicates a lower dot speed at the periphery from the FOE. A lower dot contrast indicates a weaker local motion signal. All these features simulate more challenging viewing conditions in daily life. Our study shows that FOE-tracking was most accurate when the motion signals strength for all features was high; the stimulus range that most effectively modulated intuitive tracking varied based on the stimulus feature. Saccade accuracy and smooth-tracking quality increased incrementally at each translational speed level tested, which covered the typical locomotor speed range of human observers from walking (1.2 m/s, Mohler, Thompson, Creem-Regehr, Pick & Warren, 2007), to running (2 m/s, Thorstensson & Roberthson, 1987), and cycling (4.2 m/s; Cornelissen & van den Dobbelsteen, 1999). These measures only changed when dot contrast increased from 3.6% to 8.1% but not at higher dot contrast levels, resembling previous evidence of early contrast saturation of motion processing (Allen et al., 2010; Edwards, Badcock, Nishida, 1996). The stimulus range that induced changes in eye movement measures for motion coherence was between 50% and 100%, but not at lower motion coherence levels. The higher motion coherence required to improve performance might be attributed to the short dot lifetime used in Experiment 2.

Overall, these results imply that intuitive tracking is sensitive to changes in motion signal strength, reflecting the known overlap between brain areas responsible for visual motion processing and eye movement control (e.g., middle temporal area MT/V5+, medial superior temporal area MST, ventral intra-parietal area VIP): these areas process sensory information related to visual motion (MT: e.g., Händel, Lutzenberger, Thier, & Haarmeier, 2007) and self-motion like optic flow (MST: e.g., Duffy & Wurtz, 1995; VIP: e.g., Bremmer, Duhamel, Hamed, & Graf, 2002). For instance, MT is sensitive to stimulus features like motion coherence (Händel et al., 2007), whereas neuronal responses in MST/VIP are tuned to heading directions. A majority of MSTd (dorsal) neurons prefer a radial optic flow of a FOE within 45° of straight ahead (Duffy & Wurtz, 1995). This unique profile of MST neurons has been attributed to explain why the performance of behavioral and neural decoding of heading discrimination is superior around straight ahead (Gu et al., 2010). The same areas are also involved in the control of smooth pursuit eye movements (Lisberger, 2015) and saccades to moving targets (Newsome, Wurtz, Dusteler, & Mikami, 1985). In the absence of explicit instruction or other tasks, looking at the FOE might generate a radial flow of visual information that is in line with what we experience in everyday life (where the retinal FOE is centered to straight ahead, Matthis, Muller, Bonnen, & Hayhoe, 2020).

### Intuitive tracking serves potential applications

Eye movements are often utilized as a window of perception for preverbal and non-verbal observers (e.g., preverbal infants: Dobson & Teller, 1978; non-verbal animals: Douglas, Alam, Silver, McGill, Tschetter, & Prusky, 2005). Others have combined eye movements and explicit instruction to capture visual functions (e.g., Dakin & Turnbull, 2016; Mooney, Hill, Tuzun, Alam, Carmel, & Prusky, 2018). For example, Mooney et al., (2018) adaptively reduced the contrast of a visual Gabor stimulus until the observers stopped tracking the target, producing a contrast sensitivity function in five minutes. Asking observers to track a visual target using a cursor can also reveal observers’ visual sensitivity (Bonnen, Burge, Yates, Pillow, & Cormack, 2015). However, these assessment methods still require explicit instruction given to observers about what to follow. In the current study, we quantified the benefit of explicit instruction on intuitive eye movements and showed that intuitive eye movements share characteristics with instructed eye movements, supporting the use of intuitive eye movements to study visual processing in a broader population.

These intuitive eye movements would be useful to assess visual functions in patients where verbal understanding might be limited (e.g., patients with neurocognitive disorders). For example, previous research has manipulated dot density, number of dots, and the size of the dot field to investigate spatial integration of visual motion (Warren et al., 1988; Burr et al., 1998). Manipulating these stimulus features while observers engage in intuitive tracking might reveal deficits in spatial integration. The effect of visual field defects could be investigated by restricting the stimulus area to central vs. peripheral vision, analogous to Cornelissen and van den Dobbelsteen (1999). Furthermore, aging and visual disorders such as glaucoma are associated with impairments in perceiving and discriminating the direction and speed of moving objects (Bennett, Sekuler, & Sekuler, 2007; Falkenberg & Bex, 2007; Shanidze & Verghese, 2019; Snowden & Kavanagh, 2006), especially under low contrast (Allen et al., 2010). Such impairments can impair vision-related quality of life (Roh, Selivanova, Shin, Miller, & Jackson, 2018) or driving safety (Wood, Black, Mallon, Kwan & Owsley, 2018). Given that FOE-tracking performance can be measured in only 30 seconds of viewing time, any potential assessment tool could be limited to short viewing time and therefore be applicable in a clinical setting.

Some questions remain to be addressed before the applications of intuitive tracking. First, when do observers intuitively track the FOE? Our research did not assess the situational constraints of intuitive FOE-tracking. For example, task demands might suppress FOE-tracking. Churan, von Hopffgarten, and Bremmer (2018) showed that when viewing optic flow stimulus to encode distance traveled, observers showed preferences in whether they should sample near or further away from the FOE. Combining an optic flow stimulus with a steering task, Lakshminarasimhan, Avila, Neyhart, DeAngelis, Pitkow, and Angelaki (2020) showed that observers’ eyes tracked an invisible goal for steering, rather than the FOE. In real-world eye-tracking during walking or driving, the eyes could track future landing positions of the feet (e.g., Hollands, Marple-Horvat, Henkes & Rowan, 1995; Matthis, Yate, & Hayhoe, 2018) or the inner side of the curve before a bend (e.g., Land & Lee, 1994). These studies suggest that FOE-tracking might not be reflexive. It is possible that only in the absence of other tasks, intuitive FOE-tracking occurs.

Second, where do the inter-individual differences in intuitive tracking originate, among those who track? To investigate this, researchers should establish whether inter-individual differences are reliable, and whether they could be explained by factors such as perceptual sensitivity or attention. Whereas previous research has shown a relationship between instructed tracking and psychophysical judgment of the same visual stimulus (Bonnen, Burge, Yates, Pillow, & Cormack, 2015; Mooney et al., 2018), the relation between intuitive tracking and perceptual performance remains to be tested.

## Conclusion

Many daily functions are critically dependent on our ability to perceive and quickly respond to events in complex visual scenes, such as heading direction changes induced by self-motion. Our work shows that human observers track FOE changes even when they are not explicitly instructed to do so, suggesting that this tracking behavior is intuitive. The ability to keep the eyes aligned with a shifting FOE might serve as an important gaze stabilization strategy during self-motion, facilitating the control of the body during natural locomotion, and compensating for the unstable flow experienced when the head moves rhythmically during a gait cycle (Matthis, Muller, Bonnen, & Hayhoe, 2020). Furthermore, we show that this intuitive tracking behavior shares many characteristics (e.g., overall alignment and sensitivity to motion-signal-strength manipulation) with instructed performance, supporting the use of intuitive eye movements as sensitive indicators of visual motion processing.

## Acknowledgments

This work was supported by a Michael Smith Foundation for Health Research Trainee Award and a CIHR postdoctoral fellowship to HMC, an NSERC Discovery Grant and Accelerator Supplement to MS, and a Peter Wall Institute for Advanced Studies Wall Solutions Grant awarded to MS.

## Supplementary Materials

### Effects of Time On Task

To investigate potential changes in FOE-tracking across time, we extracted alignment measures in 30-s segments in Experiment 1. **Figure S1** plots the median and individual alignment measure, showing stable performance across time segments. Statistical analysis showed no significant effect of segment on cross-correlation coefficient (*χ^2^*(4) = 1.64, *p* = .80, *W_Kendall_* = .02), time lag (*χ^2^*(4) = 0.52, *p* = .97, *W_Kendall_* = .007) or trace position error (*χ^2^*(4) = 1.89, *p* = .76, *W_Kendall_* = .025). Split-half analysis based on cross-correlation coefficient in the first segment (**Fig. S1**, above median: green dots; below median: grey circles) revealed individual differences in the effect of time: some observers who did not track in the first segment started tracking in later segments, whereas other observers who tracked in the first segment stopped tracking toward the end. Overall, we did not find evidence of time-on-task effects on FOE-eye alignment across observers, indicating that eye movement behavior was consistent over time.

**Figure S1.**
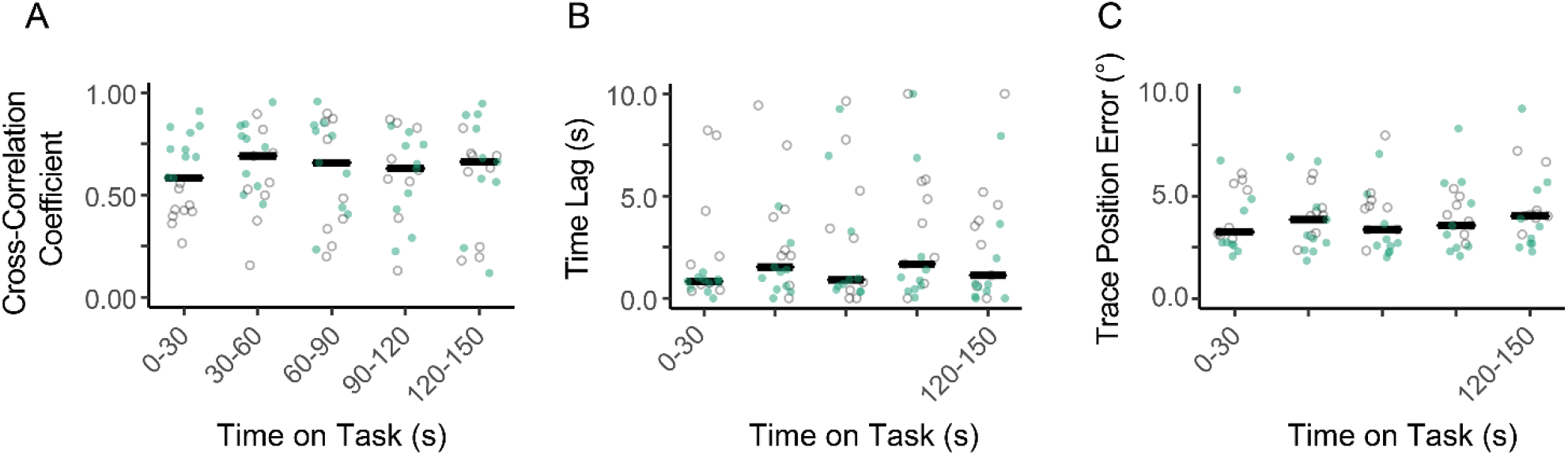
Cross-correlation coefficient (A), time lag (B), and trace position error (C) across segments of 30-s in Experiment 1. Green (solid) and grey (open) dots indicate data points from observers who had a cross-correlation coefficient in the first 30 s on-task above and below the group median, respectively. The black lines indicate the median across observers, whereas the gray dots indicate individual data points.

### Effects of Optic Flow Direction

The FOE-eye spatial and temporal alignment were similar regardless of optic flow direction in Experiment 1 (**Fig. S2**). Spatial alignment indicated by trace position error (expanding: *Mdn* = 4.0°, *SD* = 1.3°; contracting: *Mdn* = 3.9°, *SD* = 1.5°) and temporal alignment indicated by time lag (expanding: *Mdn* = 0.98 s, *SD* = 3.34 s; contracting: *Mdn* = 1.13s, *SD* = 3.35 s) did not differ significantly between optic flow directions (trace position error: *Z* = 112, *p* = .52, *r* = .12; time lag: *Z* = 96, *p* = .98, *r* = .01). Whereas the cross-correlation coefficient was smaller with expanding flow (*Mdn* = 0.59, *SD* = 0.22) vs. contracting flow (*Mdn* = 0.60, *SD* = 0.19, *Z* = 40, *p* = .026, *r* = .51), eye movement measures were superior when optic flow was expanding. For example, saccade position error was smaller with expanding flow (*Mdn* = 3.1°, *SD* =1.6°) than with contracting flow (*Mdn* = 3.9°, *SD* = 1.5°, *Z* = 146, *p* = .040, *r* = .47). Congruently, smooth-tracking velocity gain was higher with expanding flow (*Mdn* = 0.37, *SD* = 0.13) than with contracting flow (*Mdn* = 0.34, *SD* = 0.13; *Z* = 17, *p* < .001, *r* = .72). The directions of saccadic and smooth-tracking eye movements relative to the FOE were both modulated by optic flow direction. Smooth-tracking was directed further away from the FOE with expanding flow (*Mdn* = 100°, *SD* = 24°) vs. contracting flow (*Mdn* = 70°, *SD* = 17°; *Z* = 1, *p* < .001, *r* = .87), supporting the notion that smooth-tracking follows local dot motion (directed away from the FOE with expanding flow and toward the FOE with contracting flow). On the other hand, saccadic direction relative to the FOE direction was smaller with expanding flow (*Mdn* = 45°, *SD* = 21°) vs. contracting flow (*Mdn* = 55°, *SD* = 16°, *Z* = 156, *p* = .012, *r* = .56). This result does not support the notion that saccades are entirely FOE-oriented because if so, we would have seen similar saccadic direction performance regardless of optic flow direction.

Overall, tracking alignment was similar across optic flow directions. Direction analysis of eye movements showed that both saccades and smooth-tracking responded to local dot motion direction.

**Figure S2.**
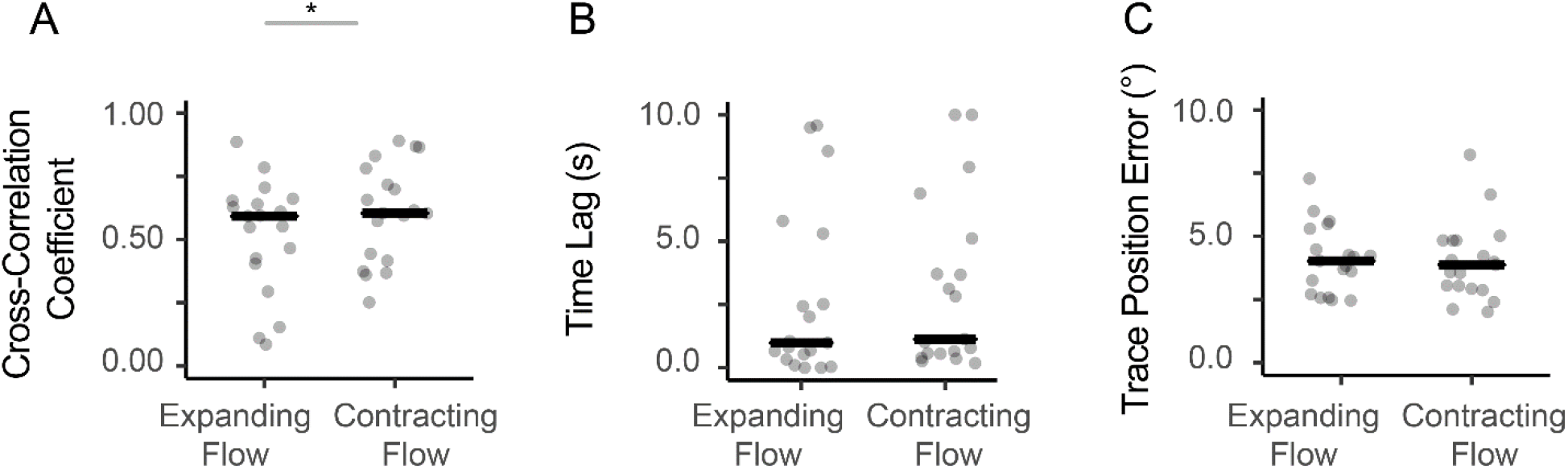
Cross-correlation coefficient (A), time lag (B), and trace position error (C) for expanding and contracting optic flow stimulus in Experiment 1. The black lines indicate the median across observers, whereas the gray dots indicate individual data points. Asterisks indicate *p*-values in pairwise comparisons (* *p* < .05).

### Interaction Between Explicit Instruction and Motion Signal Strength

Our results show that explicit instruction and motion signal strength, respectively, improved saccade accuracy and smooth-tracking quality. Here we examined the interdependency between the two manipulations by examining the effect of motion signal strength after explicit instruction and whether the effect of explicit instruction was dependent on motion signal strength. To understand the effect of motion signal strength after explicit instruction, we performed Friedman test on eye movement measures separated by each stimulus feature manipulated. Saccade position error significantly reduced when motion signal strength increased (coherence: *χ^2^*(4) = 54.1, *p* < .001, *W_Kendall_* = .62; contrast: *χ^2^*(4) = 27.8, *p* < .001, *W_Kendall_* = .33; speed: *χ^2^*(4) = 52, *p* < .001, *W_Kendall_* = .59). Similarly, smooth-tracking velocity gain was affected by motion signal strength (coherence: *χ^2^*(4) = 64.5, *p* < .001, *W_Kendall_* = .73; contrast: *χ^2^*(4) = 38.6, *p* < .001, *W_Kendall_* = .46; *χ^2^*(4) = 78.3, *p* < .001, *W_Kendall_* = .89). These effect sizes under explicit instruction were on average 75% larger than those obtained under free-viewing instruction, suggesting that changes in eye movement measures by motion signal strength were more pronounced after explicit instruction.

**Figure S3** plots the median and individual eye movement measures, contrasting free-viewing vs. explicit instruction at each motion signal strength level. To understand whether the effects of explicit instruction on eye movement measures were dependent on motion signal strength, we performed Wilcoxon signed-ranks tests comparing free-viewing vs. explicit instruction at each motion signal strength level, having adjusted for multiple comparisons. We hypothesized that explicit instruction would improve saccade accuracy when motion signal strength was strong enough for optic flow processing, but not when the signal strength was too weak to be interpreted. This hypothesis was confirmed when considering the manipulation of motion coherence. Pairwise comparisons showed that the benefit of explicit instruction on saccade accuracy was only significant when motion coherence was at or higher than 25% (*ps* ≤ .023), but not when motion coherence was low (*ps* ≥ .38). Similarly, the benefit of explicit instruction on smooth-tracking quality was only significant when motion coherence was at 25% (*p* = .005) or 50% (*p* = .017) but not at other levels (*ps* ≥ .84). Explicit instruction improved saccade accuracy at all contrast levels (*ps* < .008), whereas explicit instruction only affected smooth-tracking quality at 8.1% contrast (*p* = .011), but not the other contrast levels (*ps* ≥ .44). Explicit instruction improved saccade accuracy at all speed levels (*ps* ≤ .014) except for the fastest speed (6 m/s, *p* = .26), whereas the benefit of explicit instruction on smooth-tracking quality was significant only at the slowest speed (*p* = .003), but not at higher speed levels (*ps* ≥ .15).

Overall, we found that motion signal strength induced a larger change in eye movement measures after explicit instruction compared to free-viewing. Furthermore, explicit instruction improved saccade accuracy but not smooth-tracking quality in most tested signal strength levels, reaffirming the distinction between saccades and smooth-tracking on FOE-tracking.

**Figure S3.**
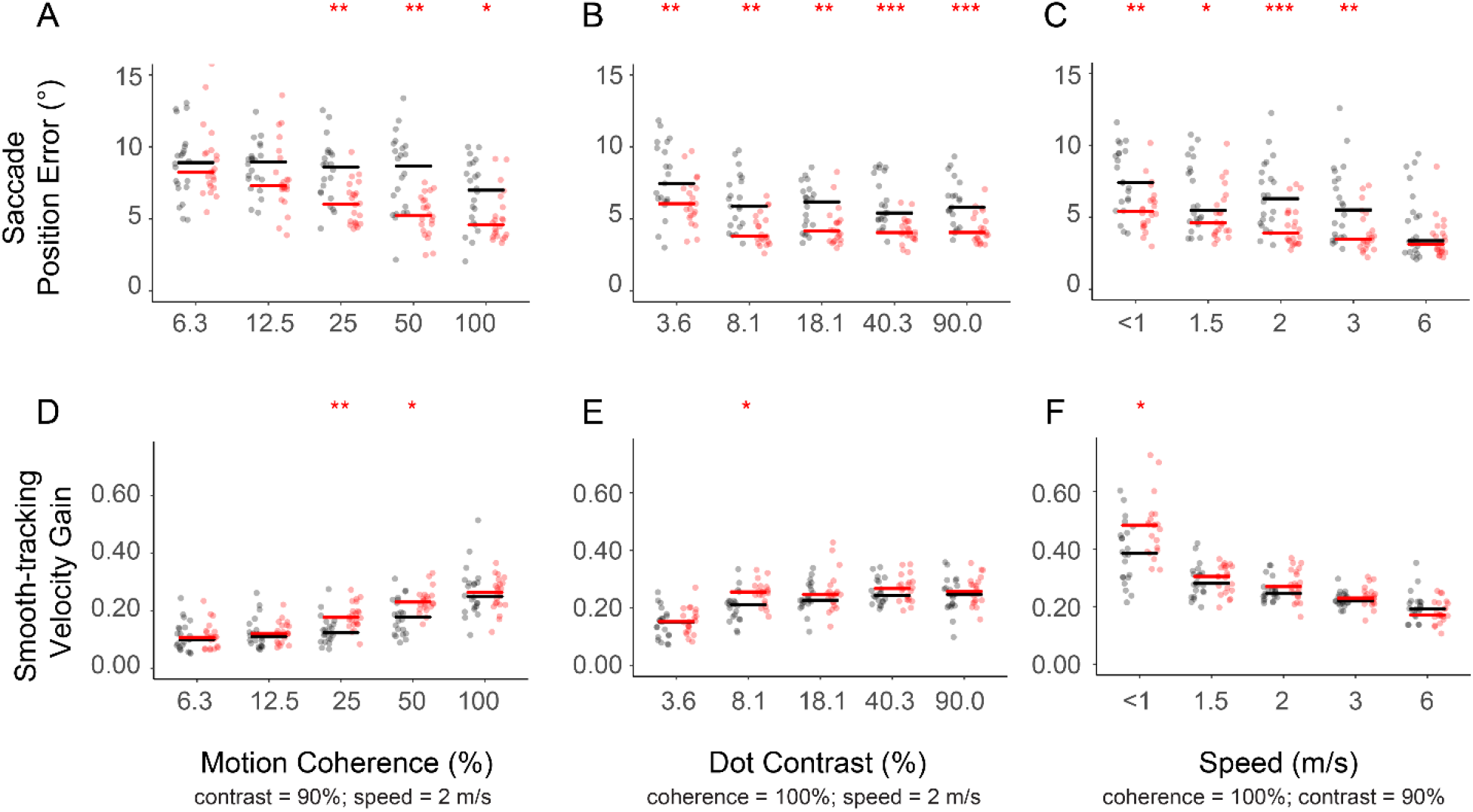
The influence of motion signal strength (horizontal axis) on saccade position error (top row) and smooth-tracking velocity gain (bottom row) in Experiment 2 under free-viewing (grey dots, black lines) and under explicit instruction (red dots, red lines), where motion coherence (A,D), dot contrast (B,E), and global translational speed (C,F) were manipulated respectively. Individual data points were plotted in colored shapes with the positions jittered along the horizontal axis to reduce overlap. The horizontal line shows the median across observers. Red asterisks indicate *p*-values in pairwise comparisons of free-viewing vs. explicit instruction (*: *p* < .05, **: *p* < .01, ***: *p* < .001).

### Citation Gender Diversity Statement

To increase the transparency of gender imbalance in reference lists in the field of neuroscience (Dworkin, Linn, Teich, Zurn, Shinohara, & Bassett, 2020), the gender balance of papers cited within this paper was quantified using a combination of automated gender-api.com estimation and manual gender determination from authors’ publicly available pronouns (see Zhou et al., 2020 for more information). Among the cited works, 59% were MM (male first-author, male last-author), 16% were WM (female first-author, male last-author), 17% were MW (male first-author, female last-author), and 8% were WW (female first-author, female last-author).

## Data Availability Statement

The data and analysis scripts used in this manuscript will be made publicly available on OSF at the time of publication.

## Notes

### Competing Interest Statement

The authors have declared no competing interest.

